# Targeting Myeloma Essential Genes using NOT Gated CAR- T Cells, a computational approach

**DOI:** 10.1101/2023.04.04.535554

**Authors:** Ieuan G Walker, James Roy, Georgina Anderson, Jose Guerrereo Lopez, Michael A Chapman

## Abstract

Sensitive cell surface proteomics studies have shown that the number of completely tumour-specific targets for adoptive cellular immunotherapy is extremely low. Even approved CAR T-cell targets appear to have expression in the central nervous system, leading to long-term neurological complications. We propose that this toxicity could be significantly improved by adoption of NOT-gates, which have been shown to limit CAR T-cell activity against healthy tissue expressing a second target that is absent on the tumour. Furthermore, the approach could also target essential, but non-specific proteins on tumour cells. The use of a NOT gate confers the specificity, whilst targeting the essential protein limits antigen escape. Here we explore the feasibility of such an approach for CAR T-cell targeting of primary myeloma. We show that none of the 45 most essential proteins are unique to the myeloma cell. However, whilst widely expressed, one of the most important proteins for myeloma cell survival, the transferrin receptor, could safely be targeted by a NOT-gate approach. Exploring co-expression patterns demonstrate 26 proteins that are not expressed on myeloma cells, but which are coexpressed with the transferrin receptor in all healthy tissues. We also describe a web app, NOTATER, which can be used by scientists with no bioinformatic capabilities to explore potential NOT-gate combinations in myeloma.

## Introduction

Multiple myeloma (MM) is a haematological malignancy, characterised by the accumulation of abnormal clonal plasma cells in the bone marrow. It accounts for approximately 10% of all haematological malignancies and is universally preceded by a monoclonal gammopathy of undetermined significance (MGUS). The hallmarks of MM include hypercalcaemia, anaemia, renal impairment, and lytic lesions in the bone which can cause pathological fractures.

In the past 15 years, MM has seen a rapid expansion in treatments: immunomodulator therapies (IMiDs; thalidomide, lenalidomide, and pomalidomide); proteasome inhibitors (bortezomib, carfilzomib, and ixazomib); and monoclonal antibody therapies targeting CD38, as well as novel antibody-drug conjugates targeting BCMA (belantumab mafodotin). These drugs are typically delivered in combination and consolidated with an autologous stem cell transplant. More recently, chimeric antigen receptor T cells (CAR T-cells) and bispecific antibodies have shown exciting therapeutic potential. However, despite this armamentarium, MM remains a fatal incurable disease.

Immunotherapies in myeloma have two main drawbacks. Firstly, MM has demonstrated several methods to escape targeting. For CD38-targeting therapies, these include down-regulation of CD38^1,2^, transcriptional changes affecting cell adhesion molecules, and CD47 upregulation to prevent phagocytosis^1^. Similar resistance mechanisms with loss of expression have been seen in BCMA-directed therapies^2^. Furthermore, BCMA can be cleaved from the cell surface by γ-secretase confirming that myeloma can persist without this molecule bound to its cell surface membrane^3^.

Secondly, toxicity of immunotherapies can be considerable, and in some cases fatal. CAR T-cells have substantial toxicity, particularly cytokine release syndrome (CRS) and immune effector cell associated neurotoxicity syndrome (ICANS). Whilst several strategies have been implemented to reduce these systemic toxicities^4^, delivery of these treatments is limited to specialised centres with expertise in managing the side effects.

Apart from CRS and ICANS, which are generic to all CAR T-cell therapies, on-target, off-tumour toxicity is a major limitation to development of novel CAR-T cells. Even with the first CAR T-cells to gain approval for myeloma therapy, which are directed against BCMA^5^ – a supposedly clean target – on-target, off-tumour toxicity has emerged as a key consideration. BCMA expression has subsequently been demonstrated in the basal ganglia and this is the likely cause of Parkinsonian-like symptoms in several patients^6^.

To address the issue of on-target, off-tumour toxicity, logic gates (referencing Boolean algebra, a mainstay of computer science and other disciplines) have been developed for CAR-T Cells. NOT-gate CAR T-cells are specifically designed to down regulate, or stop, activity when both the on-target CAR is bound together with the inhibitory CAR (iCAR)^7^. Proof of concept of NOT-GATE CAR T cells has been demonstrated in preclinical AML models^8^. Rational selection of NOT-gate combinations to target cancer currently relies on sequential trial and error, checking possible targets and inhibitory targets. In this study we leveraged our knowledge of the MM cell surface proteome to compare 965,082 possible NOT-gate CAR T cell combinations and find suitable pairs which will target MM whilst limiting off-target toxicity. We have developed this computational pipeline into a publicly available app called **NOT**-gate **A**ntigen and **T**op **E**ssential protein **R**evealer = (**NOTATER**) to help researchers rationally and safely target antigen combinations in MM.

## Methods

### Problem Formulation

For optimal selection of myeloma NOT-gated CAR T-Cells we formulated the following criteria.

1. Principal target must be expressed on the MM cell surface.
2. Principal target must have an extracellular domain.
3. Principal target must be essential to MM as defined by CRISPR screening tools.
4. Principal target ideally not expressed on T cells, to prevent fratricide during manufacturing process.
5. NOT target must not be expressed on MM cells.
6. NOT target must have an extracellular domain.
7. NOT target must always be co-ordinately expressed when the principal target is present, except on MM cells.

#### Selection of Myeloma CAR T Candidates

Primary MM cell surface proteomic data was obtained from our previous study^9^. Proteins were annotated using the Uniprot database in R^10^, and annotation for function, gene name, and extracellular length were obtained. Only proteins that contained an annotated extracellular domain were included for considerations as a target.

#### Annotation of Myeloma Essential genes

Multiple myeloma gene essentiality was obtained from Cancer Dependency Map (DepMap)^11,12^. Chronos score was used to determine gene effect^13^. Thirty-four MM cell lines were identified. Gene essentiality score was determined by the mean score across all cell lines. Only genes which were included in the cell surface protein dataset were considered.

#### Identifying Target antigen expression and NOT-gate candidates

Proteomic data was obtained from the Human Proteome Map^14^. Expression was defined as any positive expression signal detected in any tissue and was encoded as present or absent. Further annotation was performed using the Uniprot database to denote presence of an extracellular domain and known cellular location. Only proteins known to be cell surface proteins with an extracellular domain were selected as candidates for the NOT-Gate.

### Ranking of Target Antigens for Multiple Myeloma

To rank the target antigens, taking account of both expression and essentiality, potential candidates were filtered by the presence of an extracellular domain and the mean log-normalised expression of each protein was multiplied by the effect change score from the DepMap. The most negative scoring proteins were then considered the highest priority immunotherapeutic target.

#### Selection of Target proteins with NOT-Gate Partners

Using a custom script in R (version 4.2.2)^15^, NOT-gate candidates were selected from the human proteome map, such that the NOT-gate protein was expressed on all tissues that also expressed the main target antigen.

#### Development of NOTATER

NOTATER was created using the R Shiny framework and relies on dplyr (version 1.1.0)^16^ for filtering and ggplot2 (version 3.4.1)^17^ for data visualisation. This app has been made freely available for non-commercial use here: https://chapman-lab.shinyapps.io/NOTATER.

All code used in this study was complied in RStudio ^18^, and is available for review at https://github.com/ieuangw/NOTATER.

## Results

### The best MM immunotherapy targets are highly expressed on healthy tissue

We identified 777 proteins present on the myeloma cell surface membrane with an extracellular domain greater than 1 amino acid in length. To reduce the opportunity of antigen escape, we wanted to identify those proteins whose expression was most essential to the survival of MM cells. The DepMap is a large-scale publicly available data repository of multiple genone wide knock out studies. In DepMap, gene essentiality is calculated by Chronos score, which uses an algorithm to infer gene knockout fitness effects based on model of cell proliferation dynamics after CRISPR gene knockout. Genes were considered to have essential characteristics if their Chronos score was < −0.25 (**Fig. 1A**). By these criteria, 45 cell surface proteins were identified as being essential for MM survival. However, all these proteins were expressed in healthy tissue. (**Fig. 1B**).

**Figure 1.**
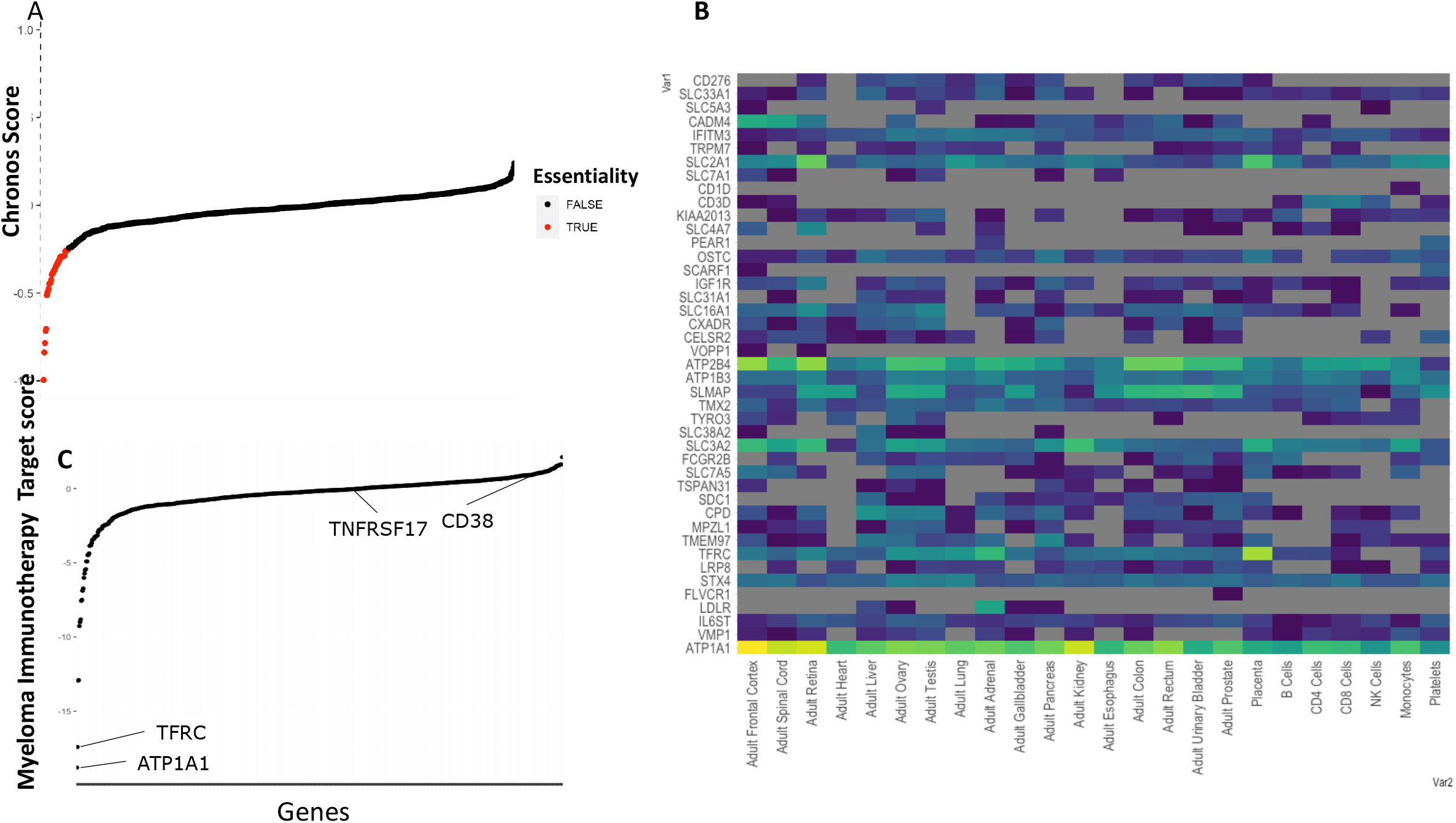
Essential multiple myeloma cell surface genes are expressed in other tissues. A) The distribution of DepMap scores of all multiple myeloma cell surface proteins ranked by essentiality. A negative score denotes a more essential gene. Genes with a score < −0.25 have been colored red and are considered essential candidates. B) Healthy tissue expression of the essential cell surface proteins highlighted in (A). (C) Re-ranking of multiple myeloma targets by a combined essentiality/expression score identifies ATP1A1 and TFRC as the top targets.

We have previously used a vector-based score to rank individual MM cell surface protein targets for single protein targeting^9^. This score accounted for both MM cell expression and off-tumour expression of the target. For a NOT-gate approach, we can discount off-tumour expression when considering the primary target. For this primary target, our major considerations were on-tumour expression in MM cells as well as essentiality of the protein for MM cell survival. We therefore combined these two features into a single score and ranked the potential targets. This approach ranked the top two targets as the cation transport ATPase, ATP1A1 and the transferrin receptor (TFRC) (**Fig. 1C**). ATP1A1 was not considered further due to the multipass nature of the protein and having a very small surface area for ScFv binding. The latter has previously been identified as a potential therapeutic target in haematological malignancies^19^. However, it was clear that both these potential targets had extensive expression outside the haematological system (**Fig. 1B**).

### There are multiple NOT-gate combinations which permit targeting of TFRC in MM

We were keen to understand the universe of NOT-gate combinations to avoid on-target, off-tumour toxicity in MM. We considered all 777 myeloma targets on the cell surface membrane, regardless of essentiality. We identified 1346 cell surface proteins in the human proteome map which were not expressed on the MM cell surface membrane. This led to 1,045,842 possible NOT-gate combinations in myeloma. Of these, 76,780 fulfilled our criteria and 4,293 combinations targeted a cell surface protein essential for MM cell survival. There were 26 iCAR candidates that could be combined with TFRC as the primary CAR T-cell target (**Fig. 2B**).

**Figure 2.**
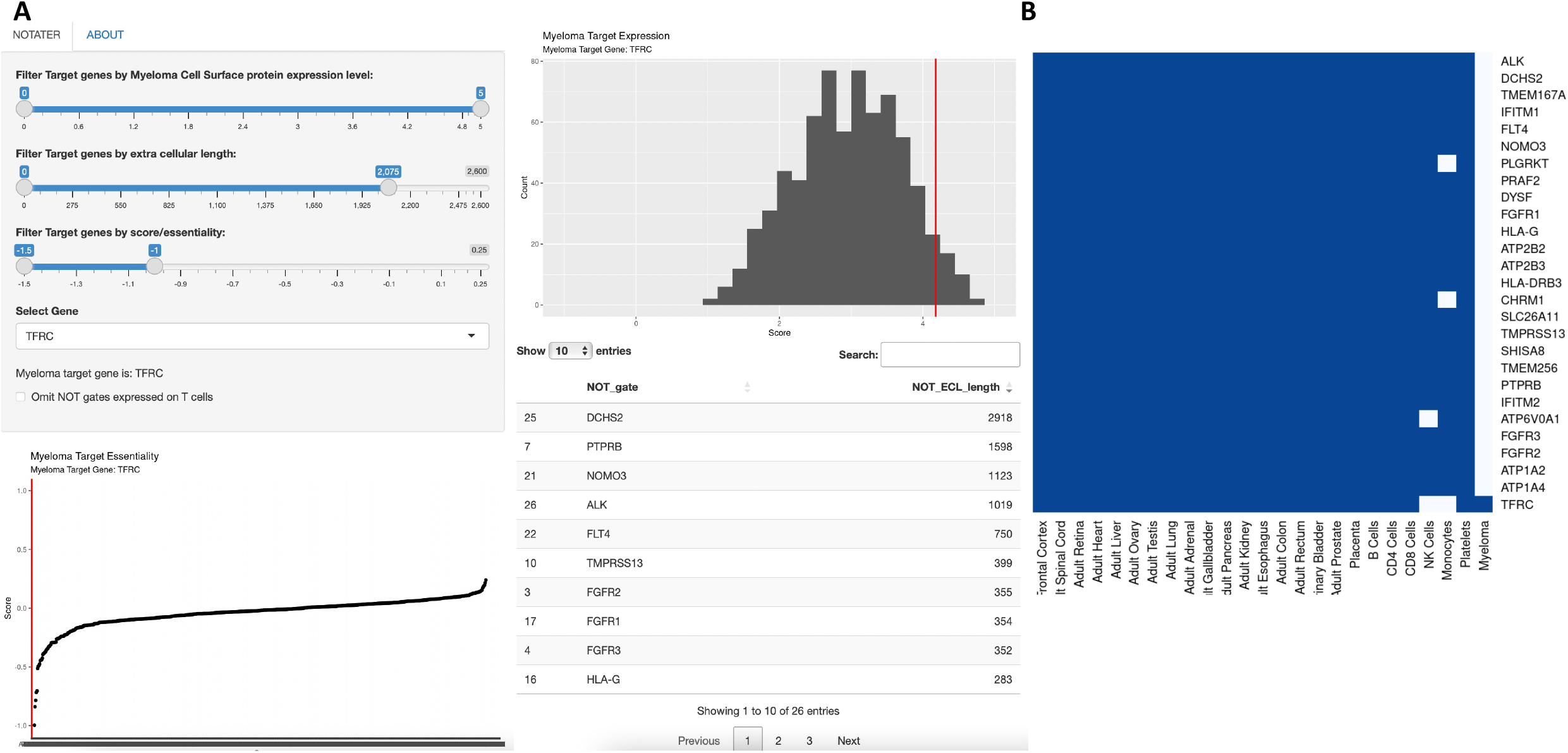
NOTATER allows selection of essential proteins and NOT-gate combinations that would be safe and effective for targeting multiple myeloma. A) NOTATER: an overview of the functionality of the app which shows how selection of combination genes for an essential gene of interest (TFRC in this case) can be chosen. B) Tissue expression of TFRC with predicted NOT-GATE combinations in a Boolean approach together with myeloma plasma membrane expression.

### Development of a broadly applicable bio-informatic tool for Multiple Myeloma tool for CARNOT Gate Combination Selection

Currently selection of NOT-Gate combinations requires bioinformatic expertise to integrate multiple datasets. To enable scientists with less bioinformatic experience to explore and assess combinations, we have developed NOTATER. This allows the user to filter Myeloma targets by their DepMap essentiality score, myeloma cell surface protein expression, and extra cellular domain size. The functionality of this can be seen in Figure 2B. NOTATER then outputs a list of proteins that could function as a NOT-gate in combination with this target. This list can be sorted by extra cellular length and filtered to remove T cell expressed targets if required. A heatmap is also displayed for users to see tissue expression of their inhibitor CAR target. NOTATER can be accessed here: https://chapman-lab.shinyapps.io/NOTATER.

## Discussion

There remains an unmet need for effective immunotherapy targets in multiple myeloma which are safe and well-tolerated. There is a growing concern about neurotoxicity with CAR T-cell therapy. Long-term neurological effects of CD19 targeted CAR T-cell therapies are starting to be reported, with suggestion of increased risk of development of epilepsy, long term memory problems^20^, and psychological disturbance^21^. More recently BCMA-targeting CAR T-Cells have shown long-term effects with persistence of movement disorders^22^. At least some of these complications are likely to be due to on-target, off-tumour toxicity. Long-term immunological side effects present an even more challenging obstacle for immunotherapy development because most target antigens are also expressed on non-malignant immune cells. Early infections within the first 12 months of CAR T-cell therapy are associated with increased mortality^23^.

Immunotherapy is currently focused on identifying a tumour-specific antigen or an antigen with minimal off-target action. However, we have previously shown^9^ that, in MM at least, there are no completely specific protein immunotherapy targets. Furthermore, basing targeting on protein specificity alone risks allowing tumours to escape immune surveillance. To reduce the risk of tumour escape, targeting genes that are predicted to be essential is a more effective strategy. Unfortunately, the most important proteins for MM cell survival have widespread off-tumour expression. To address this challenge, we have developed a rational approach to selecting CAR T-cell targets in multiple myeloma, which allows for a broader repertoire of antigens to be considered by utilising the previously published NOT-gate system^8^.

In 2022 the United States Food and Drug Administration published updated guidance for development of CAR-T cell therapy, which included advice on pre-clinical evaluation of off-target toxicity^24^. Our tool, NOTATER, allows scientists who have an interest in MM therapeutics but who do not have extensive bioinformatics experience to rapidly explore a wide number of target/inhibitor CAR combinations – thus minimizing on-target, off-tumour toxicity – prior to experimental validation. Previous attempts to develop a tool for target selection for CAR T-cells have relied on transcriptomic data^25^, which demonstrated that dual CAR T targeting outperformed single targets. However, RNA-protein correlation is poor with R^2^ values typically around 0.4^26,27^, and we have shown that, at the cell surface, the correlation is even lower. The cell surface proteomic profiling data that underpins principal target selection avoids this major confounder. The NOT-gate selection is also based on proteomic data, albeit data obtained from whole cell lysates. We have attempted to account for this by selecting proteins that have transmembrane domains, annotated extra cellular domains, and a high confidence of cellular location in the plasma membrane. Nonetheless, there remains the possibility that some potential inhibitor CAR targets are not expressed at the cell surface of off-tumour tissues, risking off targeting of healthy tissue. Therefore, confirmatory expression studies should always be undertaken prior to building CAR T-cells.

Targeting high frequency non-cancer specific antigens using the NOT-Gate approach vastly expands our target repertoire in MM. Furthermore, it allows targeting of genes previously thought to be un-targetable. TFRC has been demonstrated to be important in myeloma biology ^28,29^. A monoclonal antibody has been developed against TFRC and shown preclinical efficacy in myeloma^30^, but off tumour toxicity is expected to be considerable due to the widespread expression of TFRC on virtually all Assues. NOT-Gated CAR-T Cells represent a realistic opportunity to target these high frequency antigens while reducing this risk.

In conclusion, we present a method to integrate proteomic data with CRISPR wide knock out screens and demonstrate a bioinformatic tool which will help scientists select target combinations for NOT-gated CAR T-cells in MM.

## Notes

https://chapman-lab.shinyapps.io/NOTATER

### Competing Interest Statement

Walker- Abbvie: Consultancy, Janssen: Consultancy & Honoraria,
Chapman- Sanofi: Consultanty & Honoraria

